# Tibetan terrestrial and aquatic ecosystems collapsed with cryosphere loss inferred from sedimentary ancient metagenomics

**DOI:** 10.1101/2023.11.21.568092

**Authors:** Sisi Liu, Kathleen R. Stoof-Leichsenring, Lars Harms, Luise Schulte, Steffen Mischke, Stefan Kruse, Chengjun Zhang, Ulrike Herzschuh

**Author notes:** **Corresponding author:** Ulrike Herzschuh; Sisi Liu, **Email:**. **Author Contributions:** U.H. and S.L. conceived this study. S.L. and U.H. led the interpretation and drafted the manuscript. S.L. did the genetic lab work under the guidance of K.R.S. and L.S. S.L. and L.H. performed the bioinformatic analyses with suggestions by K.R.S., L.S. and U.H. S.L. performed the multivariate and network analyses with input by U.H. S.L. and S.K. modeled the paleoclimate change with input by U.H. S.L. simulated the spatio-temporal distribution of permafrost. S.L. performed the data visualization. S.M. and C.Z. performed the fieldwork and provided the samples. All co-authors commented on the manuscript. **Competing Interest Statement:** All authors declare no competing interests. **Classification:** Biological Sciences | Environmental Sciences.

## Abstract

Glacier and permafrost shrinkage and land-use intensification threaten diverse mountain wildlife and affect nature conservation strategy. Our understanding of alpine ecological dynamics is, however, insufficient because time series portraying ecosystem complexity adequately are missing. Here, we present an ancient metagenomic record tracing 317 terrestrial and aquatic taxa, including mammals, fish, plants, and microorganisms retrieved from a lake sediment core from the southeastern Tibetan Plateau covering the last 18,000 years. We infer that steppe-meadow turned into woodland at 14 ka (cal BP) controlled by warming-induced cryosphere loss, further driving a change of herbivore dominance from wild yak to deer. Network analyses reveal that root hemiparasitic and cushion plants are keystone taxa, likely altering the terrestrial ecosystem via facilitation. These findings refute the hypothesis of top-down control by large herbivores in the alpine ecosystem. We also find that glacier mass loss significantly contributes to considerable turnover in the aquatic community at 14 ka, transitioning from glacier-related (blue-green) algae to abundant non-glacier-preferring picocyanobacteria, macrophytes, fish, and fish-eating otters. Human impact contributes little to shaping the alpine ecosystems. By applying network analysis, we provide the first sedaDNA-based assessment of the stress-gradient hypothesis. As cryosphere loss is ongoing due to climate warming, prioritizing the protection of habitats with rich nurse plants that aid neighbors in adapting to stressful conditions is likely to be a more beneficial conservation measure than livestock reduction in the Tibetan Plateau.

**Significance statement:** Merging ancient metagenomics and network analysis gives new insights into conserving the Tibetan alpine ecosystem under ongoing warming and human perturbations. We investigated the assembly of the Yak steppe-meadow ecosystem and an alpine lake system in response to cryosphere changes over the past ∼18,000 years on the Tibetan Plateau. Large herbivores cannot be a cost-effective natural climate solution to stabilize the Tibetan alpine ecosystem because they are not keystone taxa at the ecosystem scale. Furthermore, there is no support that land use considerably shapes the alpine communities and ecosystems. Protection policy should thus prioritize focus on alpine areas with intense land use and rich in root hemiparasitic and cushion plants because these taxa act as facilitators in the ecosystem.

## Introduction

High mountain regions harbor a unique biodiversity on which human living and diverse cultures depend (1). To what extent warming, glacier retreat, permafrost thaw, and land use shape the assembly of alpine terrestrial and aquatic communities is heavily debated (2–4). Compared with other mountain areas, upland warming rates are most pronounced on the Tibetan Plateau (5). The glacier and permafrost extent (6), and thereby the world’s largest alpine ecosystem, are strongly related to temperature in this region (7, 8). Global warming threatens the unique Tibetan pastoral lifestyles (9) and biodiversity hotspots, such as the Hengduan Mountains on the southeastern Tibetan Plateau (10). Extensive cryosphere loss or even complete disappearance are predicted for the mountains of the eastern Tibetan Plateau by 2100 C.E. (11, 12). Likewise, the Tibetan Plateau lost a substantial part of its cryosphere during the last deglaciation (19−11.7 ka) (13, 14). Hence, discerning the range of ecological responses to past changes of climate, cryosphere, and land use will improve our knowledge and ability to predict future alpine ecosystem changes.

Ecological reconstructions, mainly based on pollen data, have documented a shift from alpine steppe to forest-shrub steppe/meadow on the Tibetan Plateau during the late glacial (ca. 14.7–11.7 ka) (15). Hitherto, climate change was assumed to be the main direct driver of this vegetation shift while cryosphere-driven ecological change was not considered. Also, the impact of large wild herbivores as “keystone” species and/or “top-down” engineers, such as ascribed for the Eurasian glacial mammoth steppes (16–18) has not yet been regarded as a major ecological factor for the Tibetan Plateau. Phylogenetic evidence based on modern animals indicates megafaunal migrations and population expansion on the Tibetan Plateau during the late Pleistocene by wild yak (*Bos mutus*), the largest Tibetan herbivore (19). To know whether and to what extent herbivory shaped vegetation in the past is essential for the implementation of “natural climate solutions” to vulnerable ecosystems in the future (20). However, there is no information currently available on megafaunal compositional shifts during the late glacial period on the Tibetan Plateau. It is still unknown whether changes in vegetation respond directly to changes in climate, cryosphere, or megafaunal composition.

The earliest human activities at high elevations are dated to 40–30 ka in the Nwya Devu archaeological site (4600 m above sea level/a.s.l.) on the southern Tibetan Plateau (21). Based on archaeological records, year-round habitation at high elevations (> 3500 m a.s.l.) has been widely established since 3.6 ka (22). Phylogenetic analyses infer that yak (*Bos grunniens*), the most important herding animal for Tibetans, was domesticated at 7.3 ka and population size increased six-fold between 3.6 and 0 ka (23). Similarly, pollen indicators for livestock grazing such as *Sanguisorba filiformis* and *Rumex-*type are high in records from that time (24, 25). Accordingly, there is an ongoing debate about the extent to which prehistoric land use caused the present Tibetan alpine meadow ecosystem. The mainstream perspective emphasizes that livestock grazing caused the typical Tibetan lawn vegetation (24, 26–28). The alternative viewpoint highlights the importance of abiotic drivers of past and ongoing vegetation change (29, 30). Despite the debate, land management policies aimed at restricting or removing livestock grazing have been strictly implemented at the cost of livelihoods on the Tibetan Plateau over the last twenty years (31). Moreover, since domestic herbivores and their ancestors generally share the same habitats (9), it is hard to distinguish the influence of livestock from that of wildlife grazing on vegetation composition. Accordingly, we still lack basic knowledge on the relative contribution of temperature, the cryosphere, natural herbivory, and land use on vegetation change since the last glacial period. This massively limits our ability to predict ecosystem state shifts and to provide guidance for maintaining and restoring ecological functions.

Likewise, little is known about the long-term changes of Tibetan Plateau lake communities, including macrophytes (32, 33), microbes, fish, and mammals, although the Tibetan Plateau is rich in lakes which represent unique biodiversity hotspots (34). Most studies suppose a direct response of aquatic organisms to lake-water levels (attributed to climate changes) during the late glacial period (32, 35, 36). Recently, some studies argue that glaciers, via runoff, directly contribute to a lake’s microbial composition as revealed by aquatic biomarker (n-alkanes) concentrations from Lake Hala Hu (northeastern Tibetan Plateau) (37) and microbial sedimentary DNA from Lake Yamzhog Yumco (southern Tibetan Plateau) (38). However, there is little knowledge of shifts in aquatic communities, particularly how those taxa poorly represented in the (micro-)fossil records responded to climate changes and related glacier retreats. Furthermore, during the late Holocene, human impacts on lake ecosystems are frequently inferred from a few taxa in individual assemblages such as zooplankton, algae, and submerged macrophytes. There is thus a knowledge deficit on the extent to which glacier dynamics and human-relevant activities contribute to ecosystem-level turnover in high-alpine environments.

Temporal species co-occurrence (co-existence) information is required to understand species assembly processes, including environmental filtering and biotic interactions (39, 40). According to the stress-gradient hypothesis – a major concept in ecology – positive interactions between species will become more common as environmental stress increases (41). For instance, facilitation is repeatedly reported to support the persistence of terrestrial alpine plants via microclimatic modifications from cushion plants (42, 43), a conclusion with high relevance when climate is warming (44). Those cushion plants such as *Saussurea* and *Saxifraga* are characterized by a ground-hugging mat or dense stem structure, enabling them to trap heat and soil while also providing habitable conditions for other species within their crown area. However, aside from vegetation, the stress-gradient hypothesis has been little investigated at the ecosystem level. Even more, studies supporting this hypothesis almost exclusively originate from space-for-time assessments despite species assembly being a dynamically complex process. Further, other studies from the Tibetan Plateau have shown that major ecosystem attributes such as plant richness are not analogous in space and time (45).

Lake sediments are the most suitable archives for tracking temporal species-environment relationships (46). Compared to the traditional proxies (e.g, pollen and diatoms), sedimentary ancient DNA (sedaDNA) extracted from lake sediments has become a powerful tool for retrieving more detailed assemblage of past plants, animals, and microbes within lakes and their catchments (47). SedaDNA metagenomics (shotgun sequencing) has been increasingly used for ecosystem-level investigations and shows reliable taxonomic classification down to the genus level due to advancements in bioinformatic analysis and reference databases (48, 49). To date, times-series studies of lake sedaDNA studies investigating taxa composition and co-occurrence at the ecosystem level are not available for the Tibetan Plateau.

We reconstruct terrestrial and aquatic communities by shotgun sequencing on 40 sedaDNA samples covering 17.7–0 ka extracted from Lake Naleng, located in the Hengduan Mountains, southeastern Tibetan Plateau (Fig. 1*A*). Its surrounding landscape has been influenced by Quaternary glaciation and/or permafrost (Fig. 1*A*, *B*, and *SI Appendix*, Fig. S1). The sedaDNA results reveal diverse mammals with a dominance of bovid species (e.g., wild yak) in the pre-14 ka period when steppe-meadow was well established. The Yak steppe-meadow ecosystem collapsed at 14 ka and shifted to woodlands supporting cervid species (e.g., red deer). No evidence supports large wild herbivores’ “top-down” control of vegetation shifts. Rather, warming-induced cryosphere loss directly triggered the vegetation changes that further forced the mammalian composition turnover. Likewise, we find that glacier mass loss at 14 ka strongly shaped the aquatic ecosystem from glacial microbes to a variety of interglacial organisms such as picocyanobacteria, submerged plants, fish, and otters. Overall, we infer only a limited impact of land use. After 3.6 ka, land use may have reduced mammals’ occurrences and contributed to the establishment of a picocyanobacterial turbid lake system. Through network analysis, partial correlations, controlling the influence of cryosphere changes, land use, and mediator taxa, act as surrogate representations of the direct associations among taxa. We find a high number of positive links during the glacial period (pre-14 ka, a high-stress phase) in both ecosystems, directly supporting the stress gradient hypothesis over a long time span.

**Fig. 1.**
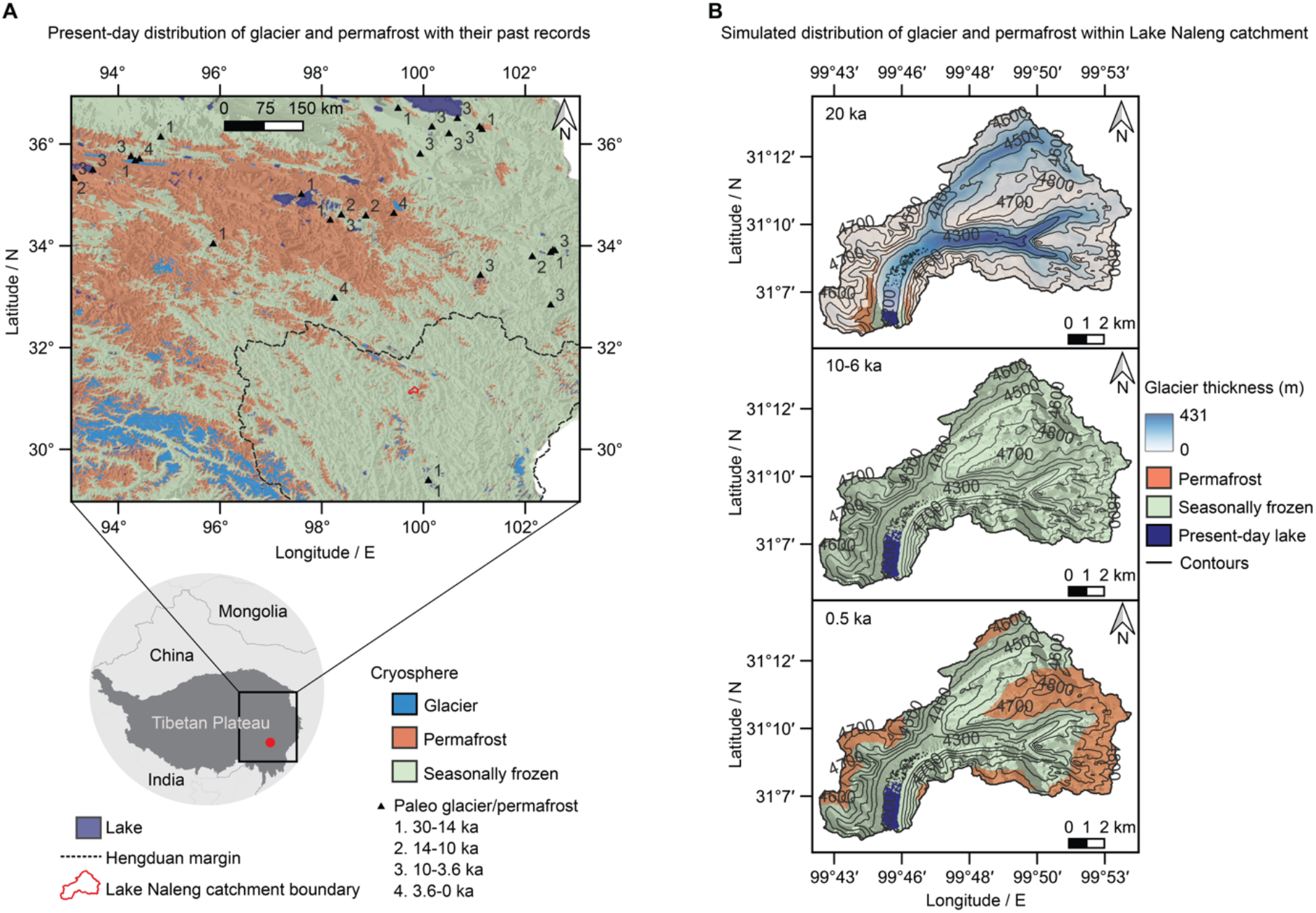
Lake Naleng, located on the southeast Tibetan Plateau (Hengduan Mountains), a global biodiversity hotspot, is influenced by the past and modern cryosphere. (A) Modern glacier and permafrost distribution (6) with their past records (14) indicate a southeastward advance of the cryosphere until 14 ka and a northeastward retreat afterward. (B) The simulated glaciers and permafrost distribution (Materials and Methods, *SI Appendix*, Fig. S1) indicate that Lake Naleng with its catchment was strongly influenced by the cryosphere during the late glacial period, while without impact during the early-to-mid Holocene. Permafrost, but not glaciers, recovered during the late Holocene, mostly in the highlands in the east of the lake catchment.

## Results and Discussion

### Ecosystem-level sedaDNA record of the past terrestrial and aquatic biosphere

Sequencing yielded 2,512,713,391 reads for bioinformatic analyses, of which 123,786,174 reads came from 40 samples and 12 controls which underwent taxonomic data cleaning and filtering (Materials and Methods, *SI Appendix*, *Supplementary Text*). Consequently, 1,067,557 reads of 317 terrestrial and aquatic taxa, comprised of seed-bearing plants, mammals, macrophytes, fish, algae, and Cyanobacteria (also called blue-green algae), with best identity ≥ 95% against the NCBI Reference Sequence Database were used for ecosystem reconstruction. The ancient origin of the reads was confirmed by read length distribution (*SI Appendix,* Fig. S2) and characteristic C-to-T substitution at the 5’ and 3’ end with sufficient reads (Materials and Methods, *SI Appendix,* Fig. S3 and Fig. S4)

The shotgun sequencing approach recovered 167 genera from 81 families of seed-bearing plants (*SI Appendix*, Datasets 1). Among these, the dominant ones are species-rich on the Tibetan Plateau and are also abundant and detected by metabarcoding and pollen analyses (25, 45, 50) from the same sedimentary core (*SI Appendix*, Table S1). The mammals, which have less biomass than plants, were traced by the shotgun sequencing approach in all samples (*SI Appendix*, Datasets 1). Most mammalian reads were identified to even-toed ungulate mammals such as Bovidae and *Cervus*, followed by *Ochotona*, a rodent-like mammal dwelling on mountains. For aquatic communities (*SI Appendix,* Datasets 2), the reads assigned to genera of aquatic macrophytes are abundant in *Potamogeton* (Potamogetonaceae) and *Myriophyllum* (Haloragaceae). Salmonidae and Cyprinidae dominated the fish assemblage, with Cyprinidae being more abundant in the high-elevation regions of the current Tibetan Plateau (51). Few reads were classified to *Lutra*, which is currently one of the endangered top predators in aquatic ecosystems (52). Most algal reads were assigned to Monodopsidaceae, specifically its genera *Nannochloropsis*, which contains species that can adapt to harsh environments such as the Last Glacial Maximum (53). As cyanobacterial blooms relate to lake ecosystem health (54), we focus on bacterial reads assigned to genera and families of Cyanobacteriota, among which *Leptolyngbya*, *Pseudanabaena*, and *Synechococcus* are the dominant genera.

Our results indicate, for the first time, that a single shotgun dataset can depict the biotic community at an ecosystem level. Our results further indicate that terrestrial and aquatic changes can concurrently be traced using the same approach. Even more, our time-series data suggest that a more continuous proxy signal can be retrieved from lake sediments compared with, for example, permafrost sediments (55). This is despite the fact that the majority of cellular organisms’ reads (98.6%) did not match any Eukaryota, which is consistent with previous environmental shotgun studies (56–58): even a small proportion of the reads (1.4%) can provide a significant amount of taxonomic information. The value of our dataset may increase in the future when more comprehensive databases are used for taxonomic assignment.

### Terrestrial ecosystem shifts from late glacial Tibetan Yak steppe-meadow to Holocene deer woodland

Overall, our results indicate an abrupt collapse of alpine steppe-meadow at 14 ka, followed by advance of subalpine shrubland during 14–3.6 ka and alpine steppe-meadow re-expansion since 3.6 ka (Fig. 2*A* and *B*).

**Fig. 2.**
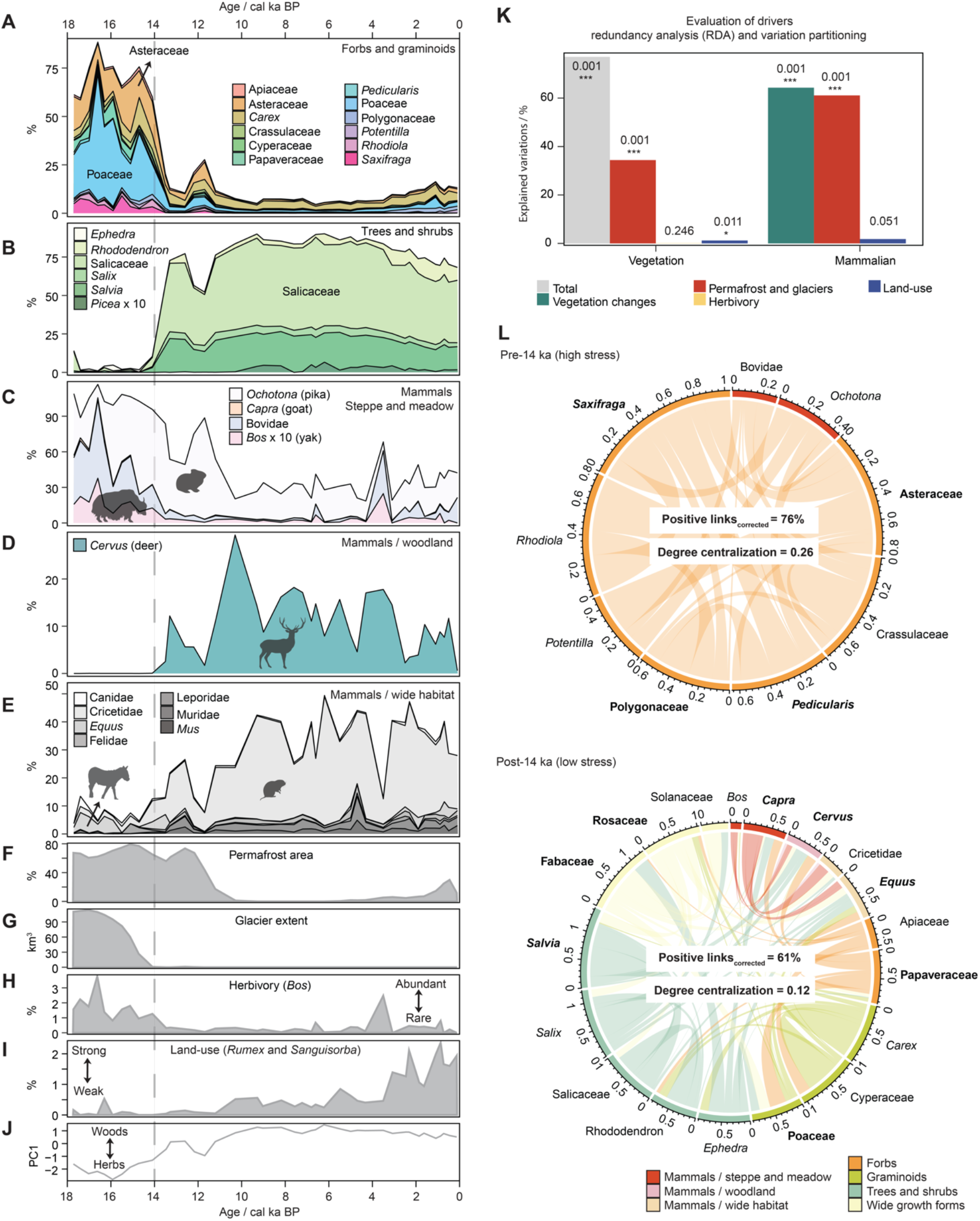
Long-term trends in shotgun-based (metagenomics) main terrestrial taxa recorded in Lake Naleng compared with temporal changes of environmental factors. (A-B) The shotgun-based relative abundance of the common terrestrial vegetation community indicates a transition from steppe-meadow pre-14 ka to woodland post-14 ka. (C-E) The shotgun-based relative abundance of the common terrestrial mammalian community shows a loss of wild megafauna since 14 ka. (F) The percentages of simulated glacier-free permafrost extent within the lake catchment (Methods and Materials). (G) The modeled glacier extent within the lake catchment (45). (H) The shotgun-based relative abundance of *Bos* relative to mammalian community as a signal of herbivory. (I) The percentages of *Rumex* and *Sanguisorba* relative to the pollen grains of terrestrial seed-bearing plants recorded in Lake Naleng as an indicator of intensity of land use (25, 50). (J) The principal curve of the terrestrial vegetation changes shows a transition from steppe-meadow with dominant herbs and graminoids in the pre-14 ka period to woodland dominated by willow shrubs and trees. (K) Evaluation of drivers based on redundancy analysis (RDA) and variation partitioning shows that cryosphere (permafrost and glacier extent) explains the highest unique portion of variation in the vegetation and mammalian communities. These results refute the perspective of top-down control by large herbivores in the ecosystem and significant land use in the creation of the modern alpine ecosystem on the Tibetan Plateau. Explained variations are represented by percentages of adjusted *r*^2^ values, obtained from RDA for joint explanation and variation partitioning for unique explanation. (L) Weighted network analyses based on positive partial correlations classify two modules, characterized by common taxa in the pre-14 ka (glacial) period with a high percentage of positive links and the post-14 ka (interglacial) period with a low percentage of positive links. Such structures support the stress-gradient hypothesis that positive interactions among organisms occur more frequently in a high stress environment. Further, more taxa connected via mediators developed in the cold phase suggesting a higher degree of centralization. Taxa with Kleinberg’s hub centrality score ≥ 0.8 (normalized score into an arbitrary range of 0–1 by taking maximum score into account) are considered keystone taxa and marked in bold. The percentage of positive links has been adjusted to the number of taxa. The chord thickness represents the positive partial correlation. The proportions of *Picea* and *Bos* are aggregated 10 times for better visibility.

The pre-14 ka vegetation is characterized by alpine forbs and graminoids including taxa dominant today in cold-dry places (e.g., *Saxifraga*, Asteraceae, and Poaceae), on moist stream banks and in meadows (e.g., *Carex*, *Pedicularis*, and Ranunculaceae) of the Tibetan highlands. During 14–3.6 ka, the abundance of woody taxa increased, with the main components being the Salicaceae taxa (e.g., *Salix*) and *Rhododendron*, as well as a portion of *Picea* during 10–3.6 ka (Fig. 2*B*). Although *Salix* is usually over-represented in sed(a)DNA spectra, its high values together with *Rhododendron* well reflect the subalpine shrub communities of the eastern Tibetan Plateau (59). Since 3.6 ka, the abundance of some forbs and graminoids (e.g., Asteraceae, Polygonaceae, and Poaceae) increased (Fig. 2*A*) even though woody taxa (e.g., Salicaceae and *Rhododendron*) were still abundant. Accordingly, the lake catchment experienced a cold-dry climate during the glacial period, followed by a moderate-to-warm and moist climate between 14 and 3.6 ka, and then returned to cold conditions afterward. The main vegetation characteristics and climate conditions agree with studies of metabarcoding and pollen analyses from the same sedimentary core (*SI Appendix,* Fig. S5*A*) as well as regional pollen records (*SI Appendix,* Fig. S5*B*).

Our shotgun data indicate that the main terrestrial mammals are medium-to large-sized ungulate herbivores (> 45 kg, Fig. 2*C* and *D*). In general, mammalian composition shifted from the steppe-meadow-adapted bovid community in the pre-14 ka period (Fig. 2*A* and *C*) to a mesic-adapted woodland cervid community post-14 ka (Fig. 2*B* and *D*). However, a compositional shift back did not co-occur with steppe-meadow re-establishment during the late Holocene (Fig. 2*A* and *C*). In the pre-14 ka period, herbivores (*Bos* and other bovids) that currently inhabit alpine ecosystems in the northern and western plateau were abundant. *Ochotona* (pika), which shares similar habitats, occurred as well (Fig. 2*C*). In the post-14 ka period, *Cervus* occurred (Fig. 2*D*); while all Bovidae species had very low proportions except for evidently increased numbers at 3.5 ka only (Fig. 2*C*). Meanwhile, our data also record a higher amount of Cricetidae sedaDNA after 14 ka (Fig. 2*E*) – which frequently occurs in the present-day moist meadows on the Tibetan Plateau. To our knowledge, this is the first sedaDNA record of the mammalian community on the Tibetan Plateau.

We investigated the key drivers of compositional changes for plants and mammals at the terrestrial community level using constrained ordination analyses (Materials and Methods). Assessed drivers include cryosphere (Fig. 2*F* and *G*, permafrost and glacier distribution simulated using temperature change and validated by sedaDNA-based permafrost and glacier microbiota changes, see Materials and Methods), herbivory (Fig. 2*H*, *Bos*%, excluded for mammalian constrained ordination), land use (Fig. 2*I*, *Rumex* and *Sanguisorba*%, ref. (24, 25)), and vegetation (Fig. 2*J*, PC1 of PCA site score, excluded for vegetation constrained ordination). All variables together explain a high amount of variation in the vegetation composition (Fig. 2*K*, 73%, *P* = 0.001). Variation partitioning (Fig. 2*K*) indicated that the cryosphere alone explains most of the vegetation variation (34.4%, *P* = 0.001) compared with the variance exclusively explained by herbivory (0.1%, *P* = 0.246) and land use (1.2%, *P* = 0.011). Such results give no support for the hypothesis of a “top-down” regulated plant community by large herbivores (16, 17). Furthermore, we find that the compositional change of mammals is strongly explained by vegetation changes (64.2%, *P* = 0.001) and cryosphere (unique 61.1%, *P* = 0.001) but poorly by land use (unique 1.8%, *P* = 0.051, Fig. 2*K*). Accordingly, the millennial-scale findings undermine the long-term argument of a human-driven ecosystem change on the Tibetan Plateau at a millennial time scale (28, 60).

An increasing number of studies have reported that co-occurrence networks can offer more information than composition on community organization, and a few studies have focused on the ecosystem level (61). We investigated species’ co-occurrence patterns at the ecosystem level (Methods and Materials) to understand direct interactions (e.g., facilitation and competition) among coexisting taxa in two different regimes, pre-14 ka (high stress) and post-14 ka (low stress). We specifically focus on positive partial correlation as an indicator of facilitation among taxa. Our aim is to assess the applicability of the stress gradient hypothesis on a millennial scale. This assessment serves as an analogy for understanding shifts in facilitation within the context of ongoing cryosphere loss and land-use changes in mountain regions.

The whole terrestrial community is clustered into two modules (Fig. 2*L*) by taking cryosphere changes and land use as predictors (*SI Appendix*, Fig. S6*A*), separating common taxa of the pre-14 ka community (e.g., Asteraceae, Bovidae, and *Ochotona*) from those dominating the post-14 ka community (e.g., *Salix*, *Cervus*, and Cricetidae). The percentage of positive links within the terrestrial community is higher in the pre-14 ka period compared with the post-14 ka interval (76% and 61%, respectively, Fig. 2*L*). This strongly supports the stress-gradient hypothesis (61) which proposes that positive interactions are promoted under stressful conditions such as the cold glacial period in our study. Furthermore, we used Kleinberg’s hub centrality score, which ranges from 0 to 1, to identify the keystone taxa, as it measures the extent of a taxon’s connections to other important taxa in the overall network (62). A taxon with a high hub centrality score is more likely to be a keystone taxon that is particularly important to the functioning of ecosystems (63, 64). We find that Bovidae and *Bos* (wild yak) show lower Kleinberg’s hub centrality scores (0.76 and 0.623) than their main food resources in their respective modules (Asteraceae: 1; Poaceae: 0.9, *SI Appendix*, Table S2), thus refuting the hypothesis of large herbivores as keystone taxa in the Tibetan terrestrial ecosystem.

A major finding is that the cryosphere governed the terrestrial Tibetan ecosystem characteristics until 14 ka by promoting a forb-dominated steppe-meadow (Fig. 2*K*) that spread in the glacier-free catchment areas on permafrost soils (Fig. 1*B* and *SI Appendix,* Fig. S1). Our results furthermore suggest that such steppe-meadow composition was “bottom-up” controlled, with plants such as *Pedicularis*, *Saxifraga* and Asteraceae species as keystone taxa, as indicated by their high Kleinberg’s hub centrality scores (Fig. 2*L* and *SI Appendix,* Table S2). *Pedicularis*, a root-hemiparasitic genus, can enhance grassland diversity by reducing the competitive advantage of grasses and legumes (which are preferred hosts) compared to forbs and sedges on the Tibetan Plateau (65). Asteraceae contain a variety of genera that form cushions (e.g., *Saussurea* and *Senecio*), which can protect diverse terrestrial alpine plants against the harsh environment on an ecosystem level by creating a favorable microclimate through their dense, low-lying cushion structure, as well as stabilizing soil, retaining water, and contributing to nutrient cycling (43, 66). Similar to *Saxifraga*, the cushion life form contributes to the diversification of other species-rich alpine genera in the Tibetan Plateau (67). These findings indicate that facilitation rather than competition shaped the glacial terrestrial Tibetan ecosystem. It represents a first assessment of the stress-gradient hypothesis over millennial time scales.

A further key finding suggested by our shotgun data is that abundant large herbivores such as wild yaks co-occurred with grass-and forb-dominated steppe-meadow (Fig. 2*A*, and *C*) extending across permafrost soils (Fig. 2*F*). This suggests that they frequently consumed protein-rich forbs rather than exclusively or heavily depending on grasses (68). The Lake Naleng’s non-pollen palynomorphs (NPP) record supports this inference, indicating high percentages of *Glomus* before 14.5 ka (69). Given their association with erosion events and the fact that their hosts, such as Asteraceae are frequently consumed by yaks in winter, an increased input of *Glomus* spores might originate from the dung of bovids after foraging on these plants (70, 71). Such consumers’ preferences are similar to the Eurasian “mammoth steppe” supporting a variety of mammals during the late Pleistocene in high latitudes (58, 72). Furthermore, vegetation shifts drive changes in mammalian communities in our study area (explained 64.2%, Fig. 2*K*), which may emphasize that resource stress (e.g., less available foraging habitats) could also pressure herbivores to facilitation rather than competition (positive partial correlations among mammals in the post-14 ka, Fig. 2*L*) which is typically less considered (73).

Overall, we suggest that temperature-driven cryosphere changes (Fig. 2*F* and *G*) controlled vegetation turnover on the Tibetan Plateau at 14 ka, which in turn promoted a decline of large herbivory. This contrasts with the proposition of megafauna as keystone engineers of the “Mammoth steppe” ecosystem of Glacial Siberia (74). Our permafrost simulation using a generalized linear model (Methods and Materials) shows that permafrost thawed at 14 ka in the U-shaped valley around Lake Naleng and along streams in response to the warming climate (Fig. 1*B* and *SI Appendix,* Fig. S1) and this is confirmed by time-series data for permafrost bacteria (*SI Appendix,* Fig. S7*A*-*C*). This likely promoted the development of alluvial soils, thereby facilitating riparian woody vegetation (*Salix* sp.) (75) as indicated in our sedaDNA record. This vegetation change may have limited the protein intake of large herbivores (e.g., yaks) (76), presumably forcing them to migrate to the restricted permafrost upslope or farther away to the northeastern Tibetan Plateau, where forbs still dominate today (15). Similarly, a decline of megafaunal grazers is reported to be coeval with dwarf willow expanding northward into the “Mammoth steppe” of northeast Siberia and North Alaska around 14.5−13.5 ka (77).

Another major result of our study suggests that cryosphere changes and not human impact (Fig. 2*K*) were the main driver of terrestrial ecosystem change on the Tibetan Plateau at a millennial time scale. Even the partial re-establishment of steppe-meadow during the late Holocene (Fig. 2*A*), which has often been related to human impact (e.g., livestock grazing, ref. (28)) could be best explained by cooling-related permafrost re-establishment in the upper Lake Naleng catchment area. Unexpectedly, a steady re-occupation by Bovidae species such as wild yak did not co-occur with the re-establishment of steppe-meadow habitats (Fig. 2*A* and *C*), nor did our record indicate any substantial herding activity post-3.6 ka. This contrasts with the co-abundance of bovid taxa and grassy alpine taxa pre-14 ka. Although crop agriculture was introduced into the southeastern Tibetan Plateau and spread up to 3600 m a.s.l. from 3.6 ka (22), cold and dry weather in this area probably forced the farmers to supplement their diets by hunting wild animals (78) and this may have contributed to a herbivore decline in contrast to the inferred high population levels associated with the pre-14 ka analog vegetation composition.

### Aquatic ecosystem shifts from a glacially impacted microbial system to a warm macrophyte fish-otter system

The aquatic ecosystem is characterized by a microbial community dominated by green algae and cyanobacteria during the cold, glaciated period before 14 ka (Fig. 3*A*), by the co-occurrences of picocyanobacteria and submerged macrophytes, fish, and otters during the warm and glacier-free early and mid-Holocene (Fig. 3*B*, *C*, and *D*), and by the dominance of non-glacially adapted picocyanobacteria after 3.6 ka (Fig. 3*B*). Evaluated drivers include glacier mass (Fig. 3*E*) and land use (Fig. 3*F*, *Rumex* and *Sanguisorba*%, ref. (24, 25)), which explain 61% (*P* = 0.001) of the variation of the aquatic communities at the ecosystem level. Variation partitioning indicates that compositional turnover is strongly related to glacier mass changes (47.5%, *P* = 0.001, Fig. 3*E*), followed by land use (6.6%, *P* = 0.001, Fig. 3*F*). These findings suggest that glacier dynamics may be the main driver of the lake’s community composition change on the Tibetan Plateau (37, 38).

**Fig. 3.**
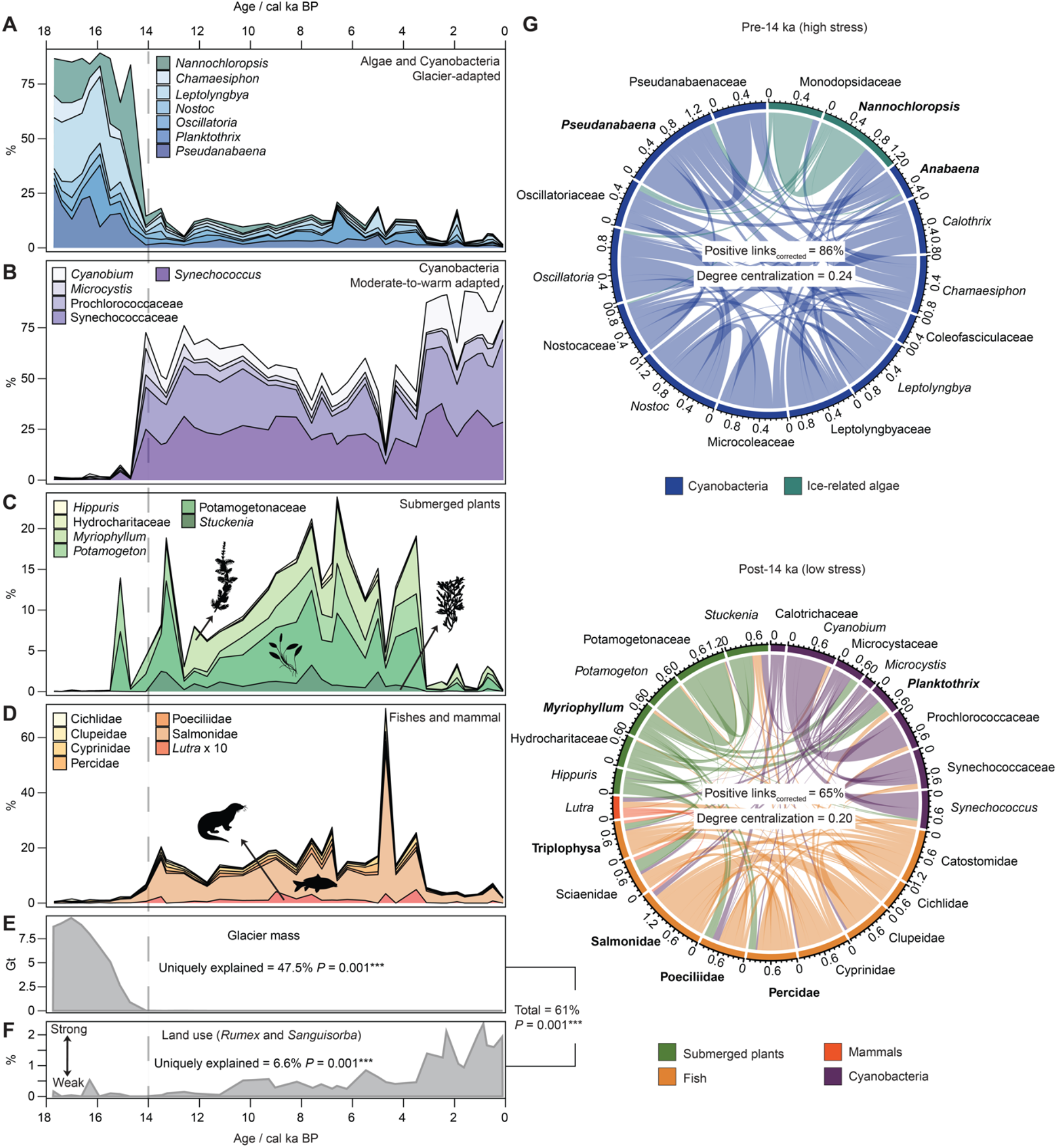
Long-term changes in shotgun-based (metagenomics) aquatic communities recorded in Lake Naleng compared with temporal changes of environmental factors. (A-D) Changes in the relative abundance of the common taxa indicate a shift from abundant glacier-adapted microbes (algae and cyanobacteria) in the pre-14 ka period to non-glacier-adapted picocyanobacteria, submerged plants, fish, and fish-eating otters until 3.6 ka, followed by overabundance of picocyanobacteria in the post-3.6 ka interval. (E-F) Evaluation of environmental factors suggests a high contribution of glacier mass on aquatic community composition changes. Explained variations are represented by adjusted *r*^2^ values, obtained from RDA for joint explanation and variation partitioning for unique explanation. (G) Weighted network analyses based on positive partial correlations classify two modules, characterized by common taxa in the pre-14 ka and post-14 ka, showing a high percentage of positive links in in the glacial period (pre-14 ka) while lower in the interglacial period (14-0 ka). Such structures support the stress-gradient hypothesis that positive interactions among organisms become prevalent under stressful environments. Further, more taxa connected via mediators developed a centralized system in the glacial phase. Taxa with Kleinberg’s hub centrality score ≥ 0.8 (normalized into an arbitrary range of 0-1 by taking the maximum score into account) are considered keystone taxa and marked in bold. The percentage of positive links has been adjusted to the number of taxa. The chord thickness represents the positive partial correlation.

Network analyzes taking glacier mass changes and land-use as predictors group aquatic taxa into two modules (Fig. 3*G*), typified by a unique community in the pre-14 ka period (e.g., *Nannochloropsis*, *Pseudanabaena*, and *Anabaena*) and post-14 ka (e.g., *Myriophyllum*, Triplophysa, Salmonidae, Poeciliidae, Percidae, and *Planktothrix*). The aquatic ecosystem exhibited a higher relative number of positive links in the pre-14 ka period (Fig. 3*G*), supporting the stress gradient hypothesis. Apart from glacier loss, land use drives partial correlations among taxa (SI Appendix, Fig. S6*B*), suggesting that Lake Naleng’s productivity and nutrient status play a role in co-occurrence pattern of the whole lake ecosystem.

Lake community inferred from shotgun data was dominated by microbial taxa, including *Nannochloropsis* (green algae), *Leptolyngbya*, and *Pseudanabaena* until 14 ka (Fig. 3*A*), when the lake catchment became glacier-free (Fig. 3*E*). These taxa or their congeners are known from cryoconite holes on a glacier’s surface in the Arctic and Asian mountains (79) and have been retrieved from Arctic lake sediments from the glacial period (53). Our results confirm that a glacier’s microbial communities are able to colonize downstream, and that glacier ecosystem changes can be traced by lake sediments (80–82). Further, the high relative abundance of *Oscillatoria*, the genera hosting rich toxin-producing strains (54, 83), in the pre-14 ka period (Fig. 3*A*) possibly forced microbes into engaging in positive associations (84, 85). Yet, whether such biotic-relevant stressors can trigger lake communities to interact positively and override the effects of abiotic stress (e.g., glacier) on millennial scales will require further evaluation.

Obviously, glacier disappearance (Fig. 3*E*) induced a transition from glacier-preferring (blue-green) algae in the pre-14 ka period to picocyanobacteria communities during post-14 ka, including the *Synechococcus* and *Cyanobium* taxa (Fig. 3*B*). Their relative abundance declined when other submerged plants such as *Potamogeton*, *Stuckenia,* and *Myriophyllum* (Fig. 3*C*) notably increased in response to warmer water conditions (86), which, in turn, favored the invasion from lower elevations and successful reproduction of fish (87, 88) including Salmonidae and Cyprinidae (Fig. 3*D*). These fish promoted the establishment of piscivores including otters (89). In addition, the submerged macrophytes can efficiently maintain water quality by releasing anti-cyanobacterial fatty acids (90). The key roles of submerged plants and fish are further inferred from co-occurrence patterns (*SI Appendix*, Table S3).

Two lake states likely coexist in one aquatic ecosystem post-14 ka, either spatially (*SI Appendix*, *Supplementary Text*) or temporally separated, i.e., a macrophyte-dominated clear-water lake system at the lake shore and a blue-green algae-dominated turbid lake system. Hence a complex subalpine lake ecosystem was established during the warm phase (post-14 ka) characterized by a low degree of centralization (Fig. 3*G*). Probably initiated by a decline in temperature-lowering submerged macrophytes, the lake ecosystem completely collapsed, leaving picocyanobacteria (*Synechococcus* and *Cyanobium* taxa, Fig. 2*B*) as the main component, which, in turn, posed a further threat to submerged plants by reducing light and oxygen (91). To what extent this change was enhanced by human impact is uncertain but it is rather unlikely as we find no clear traces of herding (e.g., livestock DNA) in the mammalian community (Fig. 2*C* and *D*). This finding highlights the importance of taking a long-term perspective when assessing the effects of natural and human-induced factors on lake state transitions, particularly before predicting the presence of critical transitions (92) and implementing sufficient lake restoration strategies to mitigate cyanobacteria blooms (54).

### Conclusions and lessons learned for risks in a warmer future and restoration if temperatures cool again

We find that climate and cryosphere-induced vegetation changes impact mammal abundance and composition, not vice versa; hence, there is no support for herbivory as a major ecosystem driver. Accordingly, managing large herbivores may not represent a conservation option; instead, only lowering the temperature by reducing global carbon emissions and preserving the cryosphere will help the conservation or restoration of the Tibetan alpine ecosystem.

We infer that *Pedicularis* and cushion plants (Asteraceae and *Saxifraga*), in particular, are keystone taxa for the Yak steppe-meadow ecosystem and should accordingly be the focus of protection measures, as they support high biodiversity at the ecosystem level by common facilitations. The way to protect them is to conserve their habitats, with priority given to areas not invaded by shrubs.

We deduce that greater grazing during the late glacial compared with the late Holocene did not destabilize the terrestrial ecosystem. By analogy, relaxed pasture management (e.g., moderate livestock reduction policy) does not represent a risk to the present Tibetan Plateau ecosystem but can be recommended to sustain contemporary livelihoods in the highlands of the Tibetan Plateau (93).

Pronounced shifts in the dominance of aquatic taxa, particularly microbial communities, occurred synchronously with the substantial decay of glaciers at 14 ka, whilst the glacial lake ecosystem was not restored in the cold late Holocene (3.6−0 ka) presumably because glaciers did not re-establish. Our findings call for more effort to inspect the glacier-lake connectivities of microbial communities as a base for developing and implementing appropriate conservation strategies. However, it is important to note that the specific aspects of glacial microbes that should be protected may vary depending on the ecosystem and the goals of conservation efforts. As our study does not investigate this aspect, we cannot offer specific recommendations for conservation actions related to glacial microbes in Tibetan glacial lakes.

Warmth-related aquatic ecosystems composed of submerged macrophytes and blue-green algae support fish and otters (main predators of fish). Warmth increased the complexity of the lake ecosystem with the coexistence of two states and less common positive associations. The lake ecosystem shifted to a turbid water state with few eukaryotes primarily due to cold-related macrophyte loss. It is unlikely that the external loading of nutrients (e.g., from husbandry) caused such a state shift as no similar signal has been observed from the terrestrial mammalian community.

We find that terrestrial and aquatic species co-occurrence patterns respond to the loss of the Tibetan cryosphere. A structure with fewer facilitative interactions suggested by positive partial correlations was detected for a permafrost-free terrestrial ecosystem and a glacier-free aquatic ecosystem. Our findings have broader implications beyond our study site, as we investigated a typical alpine lake with a catchment that includes elevational ranges typical of the southeastern Tibetan alpine ecosystem.

## Materials and Methods

### Modern site setting

Lake Naleng (31.10° N, 99.75° E; 4200 m a.s.l.) is situated in a glacier-formed basin on the southeast Tibetan Plateau, which is a biodiversity hotspot (also referred to as the Hengduan Mountains). This lake is classified as a mesotrophic lake based on its pH-value (8.11), Secchi depth (2.9 m), and dissolved oxygen content (6.86 mg/l) (measured at noon in 09.2009, ref. (69)). The catchment area is about 120 km^2^ and characterized by steep slopes and a narrow floor (U-shaped glacial trough) with Miocene granite and granodiorite rocks (94). The study area is mainly influenced by the South Asian summer monsoon and the East Asian winter monsoon. Warm and humid air masses occur in summer whilst cold and dry air masses dominate in winter. The modern catchment vegetation is alpine shrubland and meadow with two patches of forests on mountain slopes (*SI Appendix*, Supplementary Text). Human influence is livestock grazing (yaks and Tibetan sheep) in alpine meadows during the summer in the lake catchment (25).

### Late Pleistocene/Holocene site setting

The regional and local climate was generally cold and dry during the late glacial period (15), with moraines in the lake’s outlet dating to 21.5–17.5 ka (95), suggesting glaciers shaped the lake basin during and even earlier than this period. During the Holocene, a warm and humid climate has been widely recorded for the southeastern Tibetan Plateau (96) as well as our lake catchment (25, 94). These environmental settings are well captured by our cryospheric simulation (*SI Appendix*, *Supplementary Text*) and sedaDNA metagenomics (*SI Appendix*, Fig. S7*A* and *B*). Human impacts are assumed to have intensified at high elevation when the agropastoral economy expanded up to 3500 m a.s.l. around 3.6 ka (22). Based on the C/N ratios (varying between 2 and 16) and δ^13^C_org_ values (varying between – 31% and –25%) of the Naleng core (94), it is suggested that the lake sediments primarily contain the remains of aquatic organisms, and have received very little input of terrestrial plant materials through time (97).

### Core, chronology, and sedimentary ancient DNA material

The core collection, dating, and age-depth model are described in a previous study (69, 94). Due to a lack of macrofossils, sixteen samples of bulk organic carbon were dated by accelerator mass spectrometer (AMS) ^14^C at the Leibniz Institute Kiel.

The 1 cm thick sediment samples were stored at 4°C until subsampling for sedaDNA isolation that was performed using a PowerMax® Soil DNA Isolation kit (Mo Bio Laboratories, Inc. USA) with a modified protocol. A full description of the procedures is provided in Liu et al. (45).

### Library preparation and shotgun sequencing

A total of 40 sedimentary ancient DNA samples spanning 17.7–0 ka was used for library preparation. Each library batch contains sedimentary DNA isolates (15 ng), at least one DNA extraction blank, and one library blank. The libraries were prepared following the established protocol of single stranded DNA library preparation (98) with incubation of second ligation (CL53/CL72) on a thermomixer. Then, all libraries with 50 µL each were frozen at –20°C. These libraries were subjected to qualification, indexing purification, and initial quality controls (99). Four sample libraries underwent agarose gel purification to minimize the impact of adapters on targeted DNA fragments (*SI Appendix,* Supplementary Text).

Initially, libraries of 38 sediment samples and nine blanks were equimolarly pooled into four final pools of 10nM each, with a molarity ratio of 10:1 between samples and blanks. They were sequenced on a HiSeq 2500 platform (2 x 125 bp with High-Output V4 mode) and an Illumina NovaSeq 6000 (two pools on 2 x 125 bp and one pool on 2 x 150 bp), respectively. Afterward, we decided to increase the sequencing depth for 16 sedimentary libraries and sequence two additional sediments to obtain better temporal resolution during the mid-Holocene. The 18 sediment samples with four blanks were equimolarly pooled into two pools of 10nM each (with a molarity ratio as above) and sequenced on an Illumina NovaSeq 6000 (2 x 150 bp). The sequencing was performed by *Fasteris* SA (Switzerland) (*SI Appendix*, Table S4). Finally, libraries of 40 lake sediments and 12 controls yielded 631.7 GB.

### Bioinformatic analyses

We utilized FastQC (v 0.7.11, ref. (100) to assess the quality of the raw reads (2,512,713,391), and then applied clumpify (BBMAP, https://github.com/BioInfoTools/BBMap/blob/master/sh/clumpify.sh, v. 0.20.1) with default settings, except for the ‘dedupe=t’ setting, to remove duplicate raw reads. Subsequently, we used Fastp (v. 0.11.9, ref. (101), https://github.com/OpenGene/fastp) for adapter trimming and merging of paired-end reads in parallel. The deduplicated reads were trimmed for adapter filter (-a auto) when R1/R2 lacked overlaps, poly G ends (--ploy_g_min_len 10), poly X ends (–– ploy_x_min_len 10), quality filter (--qualified_quality_phred 15, –-unqualified_percent_limit 40, –-n_base_limit 5), length filter (--length_required 30), lower complexity filter (--low_complexity_filter 30). Overlapped reads were merged (-m –-merged_out) based on overlapping detection with a minimal overlapped length of 30 bp (overlap_len_require 30), a maximum mismatch limit of 5 (overlap_diff_limit 5), and a maximum percentage of mismatch of 20 (overlap_diff_percent_limit 20). Then, the outputs were evaluated by FastQC (v 0.7.11, ref. (100) for quality check.

The merged reads (1,724,066,613) were utilized for taxonomic assignment through end-to-end alignment in Bowtie2 (version 2.5.1, ref. (102)) and hereafter using ngsLCA (v. 1.0.5, ref. (49)). We established a taxonomic reference database incorporating all available RefSeq genomes from the NCBI (National Center for Biotechnology Information, downloaded on 14.08.2023). Following the HOLI pipeline (103), this database was constructed using Bowtie2 (version 2.5.1, ref. (102)) with a default setting. The merged reads were aligned against the reference genome databases to identify a maximum of 1000 valid and unique alignments (-k 1000). The resulting possible alignments were sorted and hereafter classified using ngsLCA (v. 1.0.5, ref. (49)) with a minimal identity of 95%.

### Ancient DNA (aDNA) authentication and effect of aDNA damage

Short DNA fragmentation and cytosine deamination, indicated by high frequency of C>T substitutions in the 5’ and 3’ ends (*SI Appendix*, Fig. S2 and Fig. S3), characterize our metagenomic data originating from ancient sources (104). MapDamage2 (v. 2.2.1) was applied on taxonomically classified reads with settings of –rescale and –-single-stranded (105). 26 common taxa were authenticated with sufficient reads (*SI Appendix*, Fig. S3) as representations of terrestrial mammalian community (*Bos, Cervus, Equus*, *Ochotona*, and Cricetidae), terrestrial vegetation (Poaceae, Asteraceae, *Carex*, *Rhodiola*, *Saxifraga*, *Potentilla*, *Pedicularis*, *Salix*, and *Salvia*), aquatic vegetation (*Myriophyllum and Potamogeton*), fish (Cyprinidae and Salmonidae), and aquatic microbes (*Nannochloropsis*, *Chamaesiphon*, *Leptolyngbya*, *Nostoc*, *Oscillatoria*, *Pseudanabaena*, *Planktothrix*, and *Cyanobium*).

We further observed an increase in the frequency of C>T deamination with age (*SI Appendix*, Fig. S4), indicating an effectively closed lake system that prevents DNA from being influenced by re-deposition and environmental variables (detailed discussion in *SI Appendix*, *Supplementary Text*).

### Taxonomic data cleaning and filtering

We obtained 123,786,174 reads assigned to cellular organisms, of which 107,648,162 reads classified to taxa at family or genus level (*SI Appendix*, Fig. S8*A*). These classified taxa are generally from Bacteria, followed by Eukaryotes and Archaea (*SI Appendix*, Fig. S8*B*). We detected a few proportions of reads in the blanks (in the range of 0.001 to 0.2%, *SI Appendix*, Table S5), while none of them shared similar taxa composition with the samples (*SI Appendix*, Fig. S9). Niche conservatism is evident in organisms on the Tibetan Plateau, particularly in plants adapted to harsh environmental conditions above 3000 m a.s.l. in our study region (106–108). The observed niche conservatism suggests that species within families and genera tend to preserve ancestral traits, showcasing consistent adaptive strategies for survival in challenging alpine environments over time. To detect the direct associations among taxa above species level, we kept reads belonging to natural seed-bearing plants, macrophytes, fish, and mammals (excluding Hominidae) if their families are recorded above 3000 m a.s.l. in the Tibetan Plateau. If a read is identified at the genus level but cannot be confidently matched to a known occurrence, we assigned it to the family level, assuming that the best-matched genus was absent in the taxonomic reference database. The list of referred families and genera was compiled beforehand (*SI Appendix*, Supplementary Text). Meanwhile, we kept Cyanobacteria and Monodopsidaceae (green algae) common in lake. Hereafter, we kept taxa that occur in at least two samples and have a total read count >= 5 across all samples (49). Consequently, 94% reads of our targeted taxa were retained (*SI Appendix*, Fig. S8*C*).

### Glacier and permafrost simulation

The spatial glacier extent and thickness were simulated using the numerical GC2D ice-flow model with specific settings for our study area, which have been reported in our previous study (45). The modeled glacier dynamics compare well with the changes in clay content (< 2 μm) of Lake Naleng sediments that suggest decreasing influences of glaciers after 14.5 ka (94). We further calculated the percentage of glacier extent in the lake catchment (*SI Appendix*, Fig. 10*A*) and glacier mass from 22–0 ka at 500-year intervals (*SI Appendix*, Fig. 10*B*). We interpolated permafrost distribution through time (*Appendix*, Fig. 10*C*, *D*, and *E*) from the present-day relationship between permafrost distribution and annual mean air temperature. This present-day relationship was built by a generalized linear model (GLM) and showed a significant P value < 2e-16 (t test with standard errors = 4.03e-05). We compared permafrost distribution at 0 ka with present-day to correct permafrost fields across time and assessed the reliability of the simulated results. We found simulated permafrost dynamics (*SI Appendix,* Fig. S7*A*) matching well with the relative abundance of those bacteria known from permafrost soils and active layers on the Tibetan Plateau (*SI Appendix,* Fig. S7*B* and *C*). Likewise, the simulated results are consistent with regional geological evidence (14). The detailed simulation processes and authentication are provided in *SI Appendix*, *Supplementary Text*.

### Taxonomic composition and ordination analyses

The filtered reads (1,067,557) belonging to 317 taxa were grouped into three count datasets: terrestrial vegetation, terrestrial mammalian, and aquatic ecosystem. There was a higher number of filtered reads in glacial samples than in interglacial samples for the terrestrial and aquatic datasets (*SI Appendix*, Fig. S8*C*). We calculated the relative abundance to obtain the compositionally equivalent information.

The negative binomial distributions (high mean values of read counts also have high variances) were detected in the count datasets (*SI Appendix,* Fig. S11*A*). To address heterogeneity, the count datasets were normalized using the ‘regularized log’ (rlog) transformation and the variance stabilizing transformation (vst) with default setting in the ‘DESeq2’ package (109). Both methods take sampling depth and composition into account, achieving approximately homoscedastic distributions (*SI Appendix,* Fig. S11*B* and *C*). Rlog-based counts, in particular, exhibit smaller standard deviations and were used for RDA and variation partitioning analysis.

We linearly interpolated the reconstructed past temperature (110, 111), simulated permafrost extent covering glacier-free lake catchment, and simulated glacier mass to match the temporal resolution of the shotgun data. We used principal component analysis (PCA) on rlog-based terrestrial plant counts using ‘rda(scale = FALSE)’ and extracted PC1 site scores as an indicator of terrestrial vegetation turnover. To estimate the contribution of big herbivores to terrestrial vegetation changes, we extracted the percentage of *Bos* (relative to terrestrial mammalian taxa) recorded by Lake Naleng as an herbivore intensity proxy. *Rumex* and *Sanguisorba* are considered robust indicators of land-use intensity in mountain regions (112–115) because they are commonly found in soils with high levels of nitrogen (*Rumex*) or can withstand grazing and trampling by livestock due to their basal rosette growth habit (*Sanguisorba*). Therefore, we summed up the pollen percentage of both taxa (relative to pollen grains of terrestrial seed-bearing taxa) archived in Lake Naleng as an indicator of land-use intensity.

All environmental factors were standardized using the ‘decostand(method = “standardize”)’, ensuring their transformation into variables approximating a normal distribution and achieving dimensional homogeneity (uniform unit variance). Subsequently, three rlog-transformed count datasets (terrestrial vegetation, terrestrial mammalian, and aquatic ecosystem) were separately linked to corresponding environmental variables using the ‘rda’ function. RDA, a linear method, was chosen because the length of the first ordination axis was ≤ 1 standard deviation as indicated by detrended correspondence analysis. Given temperature as a predictor variable for the cryosphere components’ simulation, the multicollinearity tests were implemented on the RDA results using the ‘vif’ function in the ‘car’ package (116) to select the independent environmental drivers (vif score ≤ 3). Then, those independent variables (*SI Appendix*, Table S6 and S7) were used to calculate the unique explanation of variation (= adjusted *r*^2^ value, ref. (117)) with variation partitioning analysis using the ‘varpart’ function.

We further verified the consistency of conclusions drawn from the ordination analyses by comparing results based on both the entire set of taxa and a subset of common taxa identified through network analysis (as explained in the next section). Both of them yielded reproducible argumentations for terrestrial vegetation, terrestrial mammalian, and aquatic ecosystem (*SI Appendix*, Table S6, Table S7, and Fig. S12*A*-*C*). These analyses were carried out using the ‘vegan’ package (118).

### Co-occurrence networks

Multiple taxa may co-occur due to responding to common environments, mediators (e.g., similar abundance changes due to shared consumers), and direct associations (e.g., facilitation) (119). To explore the direct associations, we performed a Gaussian copula graphical model to the count datasets of terrestrial (including vegetation and mammals) and aquatic. This involved using ‘stackedsdm’ to fit the marginal regression model, ‘cord’ for latent factor analysis, and ‘cgr’ for copula graph generation. These functions are available in the ecoCopula package (120). This modeling approach comprises two essential steps: (1) examining conditional dependence relationships (partial correlation obtained from the inverse of the covariance matrix) between pairs of taxa, while considering other correlations across the marginal models; (2) identifying latent drivers and discerning co-occurrence patterns resulting from direct associations from those mediated by environmental factors and mediators (120, 121). Hence, in this study, partial correlations, after accounting for the influence of cryosphere changes, land use, and mediator taxa, serve as representatives of the direct associations among taxa.

For each ecosystem, marginal regression models were established both without (H0) and with changes in cryosphere and land use (H1): H0 = stackedsdm(Y, ∼ 1 + offset(log(sizeFactors)), data = envi, family = “negative.binomial”) and H1 = stackedsdm(Y, ∼ X + offset(log(sizeFactors)), data = envi, family = “negative.binomial”). Here, Y represents the raw count data, and X represents the combination of cryosphere and land use. An offset was incorporated to adjust for variations in sampling effort, estimated using ‘estimateSizeFactors(type = “ratio”)’ in the DESeq2 package (109). Dunn-Smyth residuals and normal quantile plots suggest no violation of the assumptions of negative binomial distribution and generalized linear model for both H0 and H1 (*SI Appendix*, Fig. S13 and Fig. S14). Applying latent factor analysis using the ‘cord(nlv = 2, n.samp = 500, seed = 123)’ revealed three distinct clusters of sites based on H0, which were not evident based on H1 (*SI Appendix*, Fig. S6*A* and *B*). This suggests that misspecification in H1 has contributed minimally to the identification of direct interactions.

Subsequently, graphical models were constructed from H0 using the ‘cgr(method = “BIC”, seed = 123, n.samp = 500, n.lambda = 100)’, while for H1, the specific parameter ‘n.lambda’ was set to the optimized lambda value obtained from H0$all_graphs$lambda.opt. So, both models are comparable. To fit a completely dense graph (one margin to infer the best network), we implemented a thresholding approach, ranging from 0 to 5,000 with a step of 50, to exclude the rare taxa. The upper limit, retaining the top 5% very dominant taxa, is data-dependent and not a universal recommendation. The lowest threshold required to generate the first marginal graph was chosen for subsequent analysis.

We assumed that positive associations (e.g., facilitation) among taxa in terrestrial and aquatic ecosystems would be more prevalent during glacial periods compared to interglacial periods, in line with the stress gradient hypothesis (61). To test this assumption, we extracted the positive partial correlations from the best network of H0 and H1 for each ecosystem using ‘as_data_frame’ in the igraph package (122). The network structure was computed using ‘cluster_louvain’ with a resolution ranging from 0 to 1 in increments of 0.001. The lowest resolution value was chosen based on two criteria: (1) identical for both H0 and H1, and (2) the modules detected for each network were limited to 2.

Taken together, we determined an abundance threshold of 2,450 and a resolution of 0.69 for the terrestrial ecosystem, and an abundance threshold of 50 with a resolution of 0.52 for the aquatic ecosystem. Accordingly, 27 (694,398 reads) out of 274 terrestrial taxa (760,283 reads) and 39 (307,198 reads) out 43 aquatic taxa (307,274 reads) were used (referred to as common taxa).

We calculated five general network properties, including nodes (taxa), edges (links between taxa), degree centralization, module (community), and Kleinberg’s hub centrality scores. The positive links were adjusted as follows: the number of positive links divided by the number of full links of all taxa. We used degree centralization based on the degree of centrality scores of taxa within the module to assess how centralized the network is around a few highly connected taxa. The degree centralization score per module was calculated using ‘centr_degree(normalized = TRUE)’. High degree centralization score indicates that a few taxa within the module have significantly higher degrees (associations) than others. The loss of these taxa would significantly impact ecosystem functioning. To assess the taxon importance, we calculated Kleinberg’s hub centrality scores using “hub_score(scale = TRUE)” and considered a score of 0.8 as a threshold for being a keystone species. Such hub centrality scores are defined by the principal eigenvector of *A*\**A^T^* and rescaled between 0 and 1 by the maximum score (62).

All analyses and data visualization were performed under the R environment v.4.2.2 (123), and, unless specified otherwise, default settings were utilized.

### Data and code availability

The raw shotgun sequencing data are available in European Nucleotide Archive (ENA, www.ebi.ac.uk/ena/brower/home) with BioProject accession XXX. The source datasets to reproduce the ordination and network analyses are deposited in Dryad Digital Repository with the identifier https://doi.org/XXX. The source codes are archived at https://github.com/sisiliu-research/sedaDNA_Naleng.

## Acknowledgements

This study was funded by the Impuls-und Vernetzungsfonds − Helmholtz-Gemeinschaft (IVF, Initiative and Networking Fund of the Helmholtz Association to U.H.), Deutsche Forschungsgemeinschaft (DFG, German Research Foundation, grants 410561986 to S.K., Mi 730/1-1,2 to S.M.), and China Scholarship Council (grant 201606180048 to S.L.).

